# Cerebral bases of audiovisual temporal binding window: an awake surgery study

**DOI:** 10.64898/2026.02.03.703504

**Authors:** Solène Leblond, Robin Baurès, Tutea Atger, Marina Poinsignon, Céline Cappe, Franck-Emmanuel Roux

## Abstract

**Background:** Audiovisual integration is essential for daily functions such as speech comprehension. It relies on a temporal constraint whereby events from different sensory modalities are perceptually bound within a limited temporal window, the audiovisual temporal binding window, defining the range of stimulus onset asynchronies perceived as synchronous. While correlational neuroimaging studies (fMRI, EEG) have implicated a distributed network in audiovisual integration, the causal neural underpinnings of the temporal binding window remain largely unknown.

**Objective:** To identify cortical regions causally supporting audiovisual simultaneity judgment. *Methods:* Direct electrical stimulation (DES) was prospectively applied to 62 cortical sites during awake brain surgery in 39 patients. Patients performed an audiovisual simultaneity judgment task with varying stimuli onset asynchronies alongside standard sensory-motor, language, and visuospatial tasks. Montreal Neurological Institute coordinates were obtained for all stimulated areas.

**Results:** DES selectively impaired audiovisual simultaneity judgments while sparing other standard tasks, in 7 highly focal, right-hemispheric cortical sites (<1 cm^2^). Three sites were situated around the intraparietal sulcus, and four near the supplementary motor area. Stimulation of left-hemisphere sites produced non-selective impairments, also affecting language-related tasks.

**Conclusions:** These findings provide causal evidence for a right-lateralized frontoparietal network, involving focal regions near the intraparietal sulcus and supplementary motor area, in audiovisual temporal integration. Given the established roles of these regions in attentional and decisional processes, this study refines their contribution to the temporal binding window network and underscores the clinical importance of preserving this network during awake brain surgery.

## I) Introduction

Humans experience a constantly changing environment through multiple sensory channels, each delivering complementary information that must be integrated to create a unified perception of the external world, a process known as multisensory integration (MSI). MSI plays a crucial role in many everyday functions, notably in speech comprehension, where visual cues from lip movements facilitate the understanding [1,2]. The visual and auditory systems differ in transmission speed: although light travels faster than sound, auditory signals reach the cortex earlier than visual ones, even when both stimuli occur simultaneously [3]. Thus, the brain must flexibly integrate them to perceive synchrony while still distinguishing stimuli from separate events [4]. For auditory and visual stimuli to be integrated, they must occur within a short temporal interval, the Temporal Binding Window (TBW), so that they are perceived as synchronous [5]. This requirement is illustrated by the strong discomfort induced by even minor audio–video delays when watching a movie. To study the TBW, researchers frequently use the simultaneity judgement task (SJ).

In the SJ task, a variable stimulus-onset asynchrony (SOA) is introduced between stimuli, and participants judge whether the stimuli are synchronous or asynchronous [6–9]. Studies employing SJ tasks typically estimate the point of subjective simultaneity, defined as the SOA at which simultaneity reports are maximal. In most studies, the point of subjective simultaneity is shifted toward visual-leading SOAs (about 20–150 ms depending on the paradigm) indicating that visual signals need to precede auditory signals to be perceived as simultaneous [6,9–11]. The phenomenon is well documented in the auditory-visual perception literature [4] and is generally interpreted as reflecting the brain’s adaptation to natural sensory conditions, in which sounds typically reach the brain earlier than visual information due to faster auditory neural conduction.

Some studies showed MSI processes could rely on a broad and interconnected network [12]. Crossmodal interactions can be observed at very early stages of sensory processing, including primary auditory and visual cortices [13]. Anatomical tracing studies have revealed direct projections between early sensory areas, suggesting that crossmodal communication does not rely exclusively on higher-order convergence zones [14–17]. Physiological evidence further supports this view, as neuronal populations in both primary auditory and visual cortices exhibit modulatory responses to inputs from other modalities [18–20]. Beyond these early interactions, higher-order regions including the prefrontal and parietal cortices could play a crucial role by exerting top-down modulation to integrate sensory inputs into a unified percept [21–23]. The superior temporal sulcus (STS) has also emerged as a hub region for MSI, consistently identified across neuroimaging, electrophysiological, and stimulation studies [24–27]. A study compared brain activation during an SJ task and an attention-to-orientation task using fMRI and showed differential activation of a network, mainly in the left hemisphere, including the left anterior superior temporal gyrus, the left inferior parietal cortex, the left medial frontal gyrus, and the right operculum [28]. However, a review reported no clear lateralization and identified a broad brain network responsible for the detection of asynchrony, including, in both hemispheres, the auditory and visual cortices, the dorsal fronto-parietal attention network, the superior colliculi, the insula, the inferior parietal cortex, and the STS [29]. Consistent with this view, the neural basis of simultaneity judgment has been shown to rely primarily on a broad fronto-parietal network, which is also involved in temporal processing and selective attention [30]. Taken together, these studies converge on the conclusion that the fronto-parietal network, particularly the inferior parietal cortex, in interaction with temporal and sensory cortices, plays a crucial role in the TBW. At the same time, additional structures such as the superior colliculi and insula appear to contribute more variably depending on task demands and perceptual context. However, the presence of potential hemispheric lateralization in these areas remains debated.

In these studies, the methods used (EEG, fMRI) were correlational. However, correlational approaches alone cannot establish a direct causal relationship between the function being tested and the brain area involved. The importance of combining three types of approaches, correlational, lesional, and causal methods, for effective brain mapping has been emphasized [31].

The goal of the present study was to investigate the neural mechanisms underlying the detection of auditory-visual asynchrony. Participants performed an SJ task using only auditory-first pairs to facilitate task performance. We adopted a causal approach with maximal spatial and temporal resolution, based on the methodology of Baurès et al., employing direct electrical stimulation (DES) during awake brain surgery [32,33]. Disruptions in simultaneity judgments during localized cortical stimulation, relative to non-stimulated trials, were interpreted as evidence that the stimulated area contributed to the perception of AV synchrony.

## II) Methods

### 1. Ethics and participants

The study received ethical approval from a French *Comité de Protection des Personnes* (CPP), under authorization number NCT04128306 (RC31/18/0240 - Toulouse University Hospital). A total of 55 patients, including 18 women, all diagnosed with a brain tumour, were included during 4 years. However, 7 patients were excluded from the analysis due to their inability to perform the task during the preoperative testing, and 9 during their surgery. Therefore, the final sample on which the presented results are based consists of 39 patients (mean age: 50 ± 17 years, 18–70 years), including 13 women. Exclusion criteria included: patients under 18 years of age, individuals with aphasia or a neglect syndrome, and those unable to comprehend or perform the required tasks.

### 2. Simultaneity Judgment (SJ) Task protocol

The SJ task was administered using a Dell laptop equipped with a 1.6 GHz i7 processor and a 13.3-inch display (resolution: 1920 × 1080 pixels; dimensions: 29.5 × 17 cm, horizontal x vertical). The task was programmed and executed using EventIDE software.

Patients performed a simultaneity judgment task in which they were presented with an auditory and a visual stimulus. The auditory stimulus consisted of a pure tone at 2000 Hz and lasting 50 ms, delivered through the computer’s built-in speakers, positioned approximately 0.5 meter from the patient, with a sound intensity of 90 dB. The visual stimulus was a white circle and was displayed for 50 ms at the center of a black screen. A variable delay, referred to as the stimulus onset asynchrony (SOA), was introduced between the onsets of the auditory and visual stimuli. For each trial, patients were asked to indicate whether they perceived the sound and the visual stimulus as perfectly synchronous (**Figure 1**).

**Figure 1.**
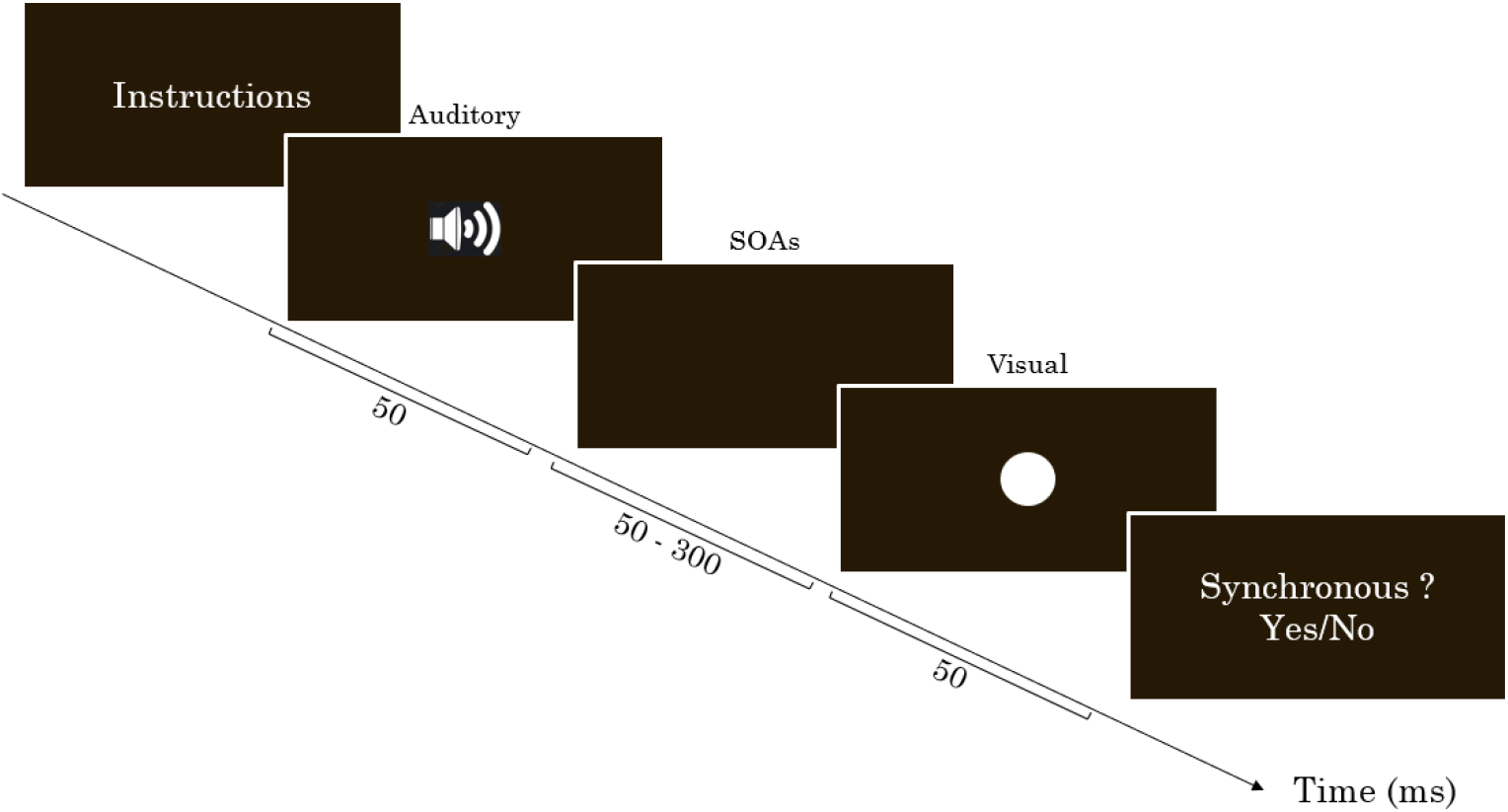
Trial structure of the Simultaneity Judgment (SJ) Task, illustrating AV stimulus presentation and patient response.

### 3. SJ task preoperative tests

Patients performed a preoperative session of the task the day before surgery to determine their individual performance level. The SOA between the auditory and visual stimuli ranged from 0 to 350 ms, with 15 possible SOAs separated by 25 ms each. Each patient completed 4 blocks of 45 randomized trials (15 SOA × 3 repetitions), for a total of 180 trials. These preoperative data were analyzed using R for each patient. For each patient, the r^2^ and slope of the linear regression curve, representing the percentage of ‘asynchronous’ responses as a function of the SOA, were extracted. The mean and 95% confidence interval (CI_95_) of these parameters were then calculated across patients. This curve allowed the identification of individual SOA values corresponding to 10% (SOA_10_) and 90% (SOA_90_) of ‘synchronous’ responses, which were subsequently used for intraoperative testing.

### 4. SJ task intraoperative tests: direct electrical stimulation

The main advantage of this brain mapping technique is its accuracy. This stimulation is expected to transiently deactivate a small cortical area, less than 5 mm × 5 mm [34,35]. Our awake brain mapping protocol was developed based on 25 years of experience [35] and used successfully following the methodology described in the studies of Baurès et al. [32,33]. Cortical stimulation was applied with a bipolar electrode (1 mm contacts) delivering biphasic square-wave pulses of 1 ms duration in a 50 Hz train, with a maximum train duration of 5 seconds. To avoid primary sensory-motor responses in the analysis of the data, pre- and postcentral gyrus responses were excluded from this study.

Depending on tumour localisation and usefulness, each cortical area tested with the SJ task was also tested with at least another task: a language task (object naming) [35], a line bisection test assessing visuospatial attention performance [36], or sensorimotor tasks. Because of clinical constraints, we tested each patient in 1 to 3 areas with the SJ task. During surgery, the exact positive stimulation areas were identified using neuronavigation (BrainLab) and registered onto the patient-specific 3D images.

Pre- and intraoperative SJ tasks were similar during operations. Nevertheless, due to the more constrained surgical context, the number of trials had to be limited. Therefore, each patient was tested using only the two individually determined SOA values from the preoperative session: SOA_10_ and SOA_90_. For each cortical area tested with direct electrical stimulation (DES), a total of 12 trials were conducted: 6 with the SOA_10_ and 6 with the SOA_90_. For each trial, the neurosurgeon applied stimulation to the selected area, and an experimenter initiated the trial approximately 1-3 s after stimulation. A break of 20 s was made between two consecutive trials. Each patient was also tested with ‘placebo’ trials, during which no actual stimulation was delivered to control for potential effects related to the surgical context. Patients unable to complete these placebo trials were excluded from the study.

Post-operative statistical analysis (***detailed in supplementary material 1***) of SJ data defined three possible outcomes:

1. If the patient’s responses remained consistent with pre-surgery performance (i.e., mostly ‘asynchronous’ for SOA_10_ and mostly ‘synchronous’ for SOA_90_), both with stimulation and during placebo trials, the area was identified as non-engaged in the SJ task.
2. If the patient showed altered performances during stimulation trials (e.g., reporting ‘synchronous’ too often at SOA_10_ or ‘asynchronous’ too often at SOA_90_), the area was considered involved in the SJ task. If the stimulation also led to language, motor, or visuospatial attention interference, we qualified the area as a ‘SJ nonspecific area,’ indicating that the synchrony perception was not the only function impaired by the stimulation.
3. If stimulation affected only the SJ task without interfering with language, motor, or visuospatial functions, the area was identified as ‘SJ specific area’. However, we cannot exclude the possibility that this ‘specific’ area may affect other untested functions.

All cortical areas were mapped on 3D cortical surface reconstructions of either the left or right hemisphere of one individual brain (case 12) included in the PALS (population-average, landmark- and surface-based) atlas [37], using the Caret software [38] and normalized to MNI space. The MNI coordinates (X, Y, Z) for each positive stimulation area were recorded (***see supplementary material 2* and 3** for coordinates, effect on the SJ task, and potential interference with other tasks).

## III) Results

### 1. Preoperative results

A linear regression of the percentage of ‘synchronous’ responses as a function of SOA was then calculated for each patient during preoperative testing (see **Figure 2** for an example).

**Figure 2.**
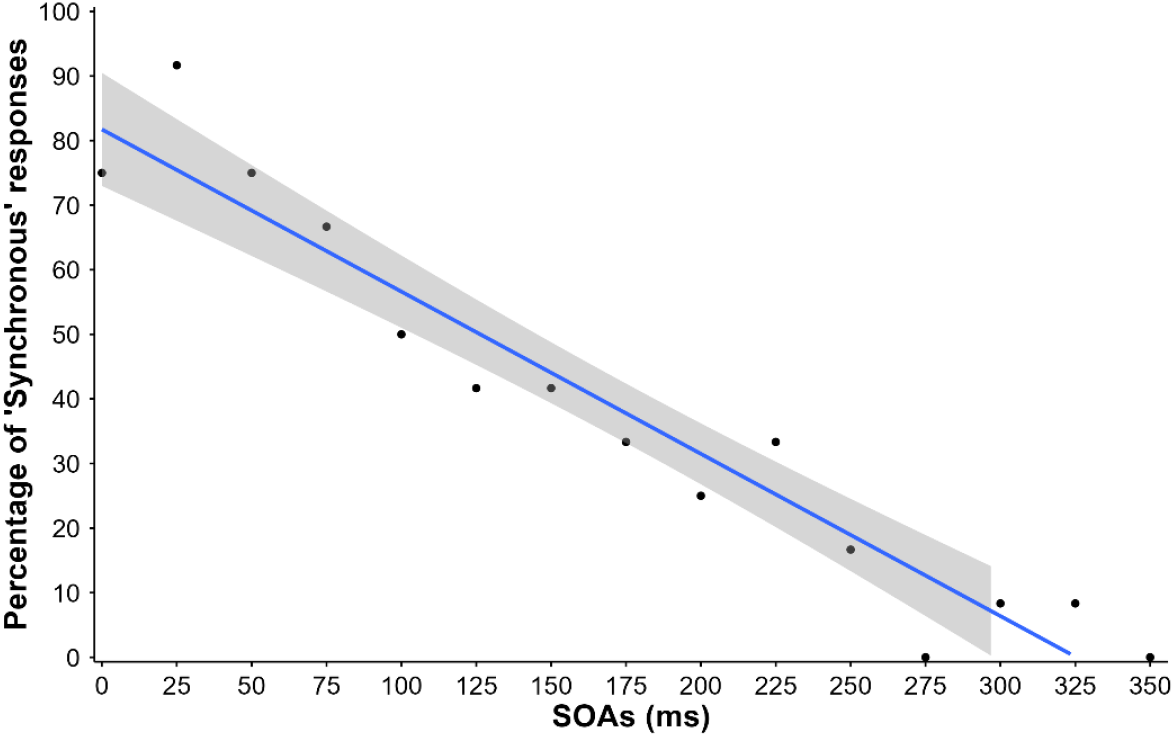
Preoperative test results for one representative patient. The blue line shows the linear regression of the percentage of ‘synchronous’ responses across SOAs, with the shaded grey area representing the 95% confidence interval (CI95).

Participants’ regressions had a mean r^2^ = .79, 95% CI = [0.76: 0.82]. The mean slope of the regression lines was –0.28, 95% CI = [–0.30: –0.26]. The SOA values corresponding to the 10% and 90% points of ‘synchronous’ responses were also calculated, as they would be used during intraoperative tests. The mean SOA_10_ was 346 ms, 95% CI = [323: 370], and the mean SOA_90_ was 102 ms, 95% CI = [93: 112].

### 2. Intraoperative results

The 62 discrete cortical areas stimulated were classified into three categories (**Figure 3**):

**Figure 3.**
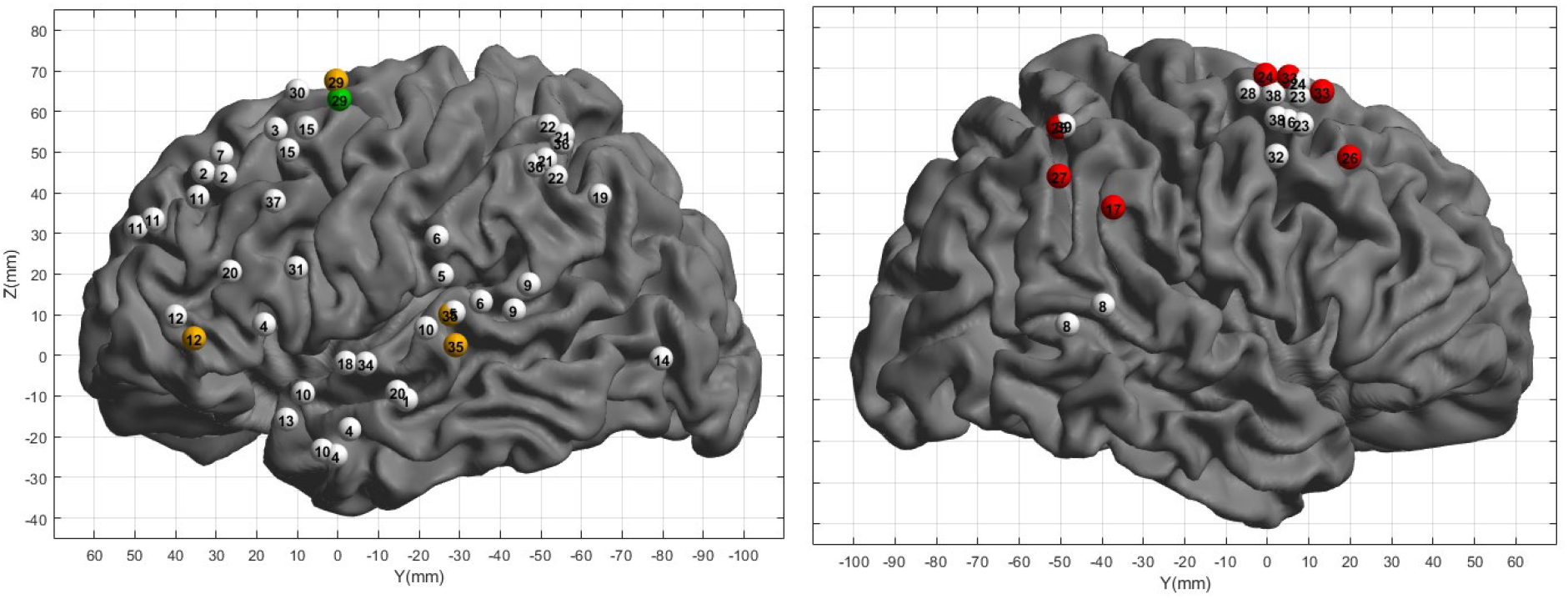
Localization of positive areas on the left and right hemispheres (MNI coordinates – **see detailed tables in supplementary material 2 and 3**).

1. Non-engaged in the SJ task (white dots), where stimulation did not alter performance on the SJ task.
2. SJ-nonspecific areas where stimulation impaired SJ task performance in conjunction with either oculomotor disturbances (green dot) or language disruptions (orange dots).
3. SJ-specific areas (red dots), where stimulation selectively disrupted performance on the SJ task without affecting any of the other tested tasks. These positive sites had sharp borders, such that a slight displacement of the electrode, as little as 5 mm within the same gyrus, abolished the interference. A substantial degree of inter-individual variability was observed, which is commonly reported when stimulating non-primary cortices in humans.

Of the 62 stimulated sites, 50 did not disrupt SJ task performance (white points). These white points indicate areas not involved in the SJ task, but that may be implicated in other cognitive or sensorimotor functions. Five stimulation sites were classified as SJ-nonspecific areas, as they induced disruptions in the SJ task along with impairments in other functions: one site (in green) located in the left superior frontal sulcus (patient 29) elicited oculomotor disturbances, and four sites (in orange) interfered with language processing (all in left hemispheres: pars triangularis (patient 12), superior and middle temporal gyri (patient 35), and supplementary moto area (patient 29)). Over the 7 SJ-specific areas (in red), 3 were located in the right supramarginal gyrus (patient 17) and the right intraparietal sulcus (patients 25 and 27), and 4 were within the right frontal lobe, particularly in the supplementary motor area (Patients 24 and 33) and the middle frontal gyrus (Patient 26). No stimulation site eliciting neglect was detected in this series.

## IV) Discussion

The study first supports a right-hemispheric lateralization of the audiovisual synchrony perception, with critical sites located near the intraparietal sulcus and the supplementary motor area (in red, **Figure 4**). We also found that synchrony perception relied on highly localized cortical patches, characterized by sharp functional borders with adjacent cortex and substantial inter-individual variability. Finally, impairments of audiovisual synchrony judgments in some left-hemispheric sites that were also involved in language tasks suggest a shared, downstream processing stage between these two functions, likely related to attentional control, as proposed in previous work [33].

**Figure 4.**
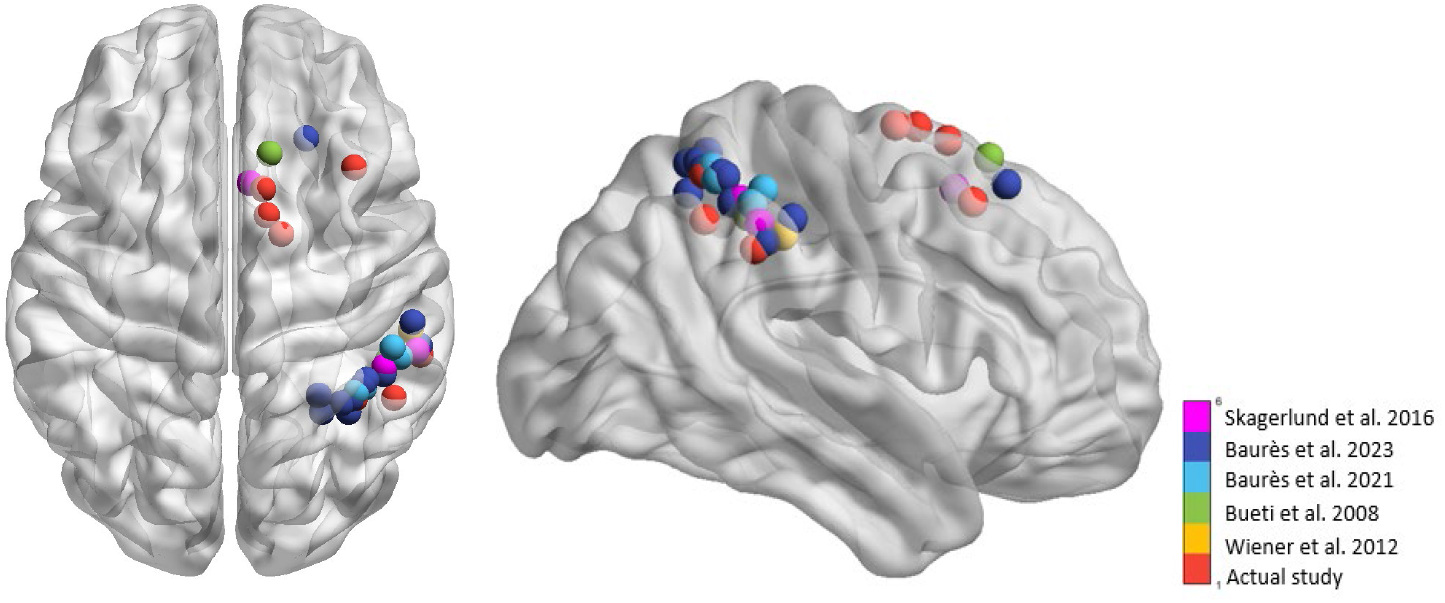
Localization of areas involved in the temporal judgment task projected onto a normalized brain using BrainNet Viewer based on MNI coordinates. Areas from the present DES study are shown in red; areas from Baurès et al. (2021, 2023) in light and dark blue; the right supramarginal gyrus area from Wiener et al. (2010, 2012) in yellow; and from Bueti et al. (2008) in green; and the right intraparietal sulcus and pre-SMA areas from Skagerlund et al. (2016) in violet.

These findings converge with evidence implicating posterior parietal regions in temporal processing. For instance, transcranial direct current stimulation during an SJ task showed that modulation of the right posterior parietal cortex alters audiovisual temporal judgments, pointing to a right-lateralized contribution to the TBW [39]. Consistent with this, Wiener et al. used rTMS and combined rTMS–EEG and demonstrated that stimulation of the right supramarginal gyrus increased the perceived duration of visual stimuli, further supporting the idea that this region contributes to temporal computations relevant for distinguishing synchronous from asynchronous events [40,41]. The supramarginal gyrus site identified in the present DES study lies very close to the coordinates reported by Wiener et al. (yellow point in **Figure 4**) [41], and to a right supramarginal region implicated in temporal reproduction in Bueti et al. (green point in **Figure 4**) [42], reinforcing the notion that this inferior parietal region is a key node within the network underlying temporal processing.

Our results also reveal SJ-specific areas within the frontal lobe, particularly in the right supplementary motor area and the right middle frontal gyrus (in red, **Figure 4**). This finding is consistent with the frequent implication of the supplementary motor area in temporal estimation and timing-specific judgments [42–44]. These results also agree with findings identifying a vast network involved in the TBW, including primary visual and auditory areas, the superior temporal sulcus (STS), and frontal regions such as motor areas and prefrontal cortex, with two-way communication between these areas [29].

The literature appears divided regarding a possible specific hemispheric lateralization of temporal processing. For example, some fMRI studies have emphasized left posterior parietal involvement during temporal-order or SJ tasks [45,46], whereas transcranial direct current stimulation results have demonstrated only a role for the right posterior parietal cortex [39]. In a literature review arguing against hemisphere specialization it was found that no right-sided lateralization was present; instead, a network involving both hemispheres equally was identified, a pattern also supported by subsequent research [29,30]. In our study, all SJ-specific sites were localized in the right hemisphere. The left hemisphere was extensively investigated with 44 stimulated sites (N=26). Five sites produced stimulation-related impairments in the SJ task, but all were non-specific: one site also elicited motor effects, and four sites were associated with language errors. These areas may contribute to the SJ task, but our protocol may not have revealed a specific role. It is also possible that stimulation of these areas altered attentional processes common to both language and TBW.

fMRI studies have consistently implicated the STS in synchrony perception [47–49], identifying it as a central hub for multisensory integration involved in simultaneity perception [29]. However, in our study, the STS was not directly involved in either hemisphere: it was only marginally explored on the right due to the tumour location, and left-hemisphere stimulation mostly affected nearby language-related cortices. Methodological differences likely contribute to this discrepancy, as DES provides causal evidence based on focal, transient disruption. In contrast, most studies implicating the STS in TBW rely on correlational measures and cannot establish a direct causal link with SJ performance.

By virtue of its spatial accuracy, DES allows dissociating the cognitive functions supported by small, neighboring cortical areas, even within highly specific cognitive or language domains. This ‘mosaic’, patchy organization is a standard observation in many electrostimulation studies [50–52]. The present study shows that the neural bases of audiovisual synchrony perception involve, with a certain degree of inter-individual variability, a core of discrete, highly localized cortical sites that may act as relays within a network initiated in the primary auditory and visual areas.

This organization is similar to findings from awake-surgery studies investigating the ‘time-to-contact’ process, defined as the remaining time before a moving object reaches a target [32,33], which was tested in a purely visual condition. Their work similarly supported right-hemispheric lateralization and revealed a specific involvement of regions surrounding the intraparietal sulcus, closely matching the parietal areas identified here (light and dark blue points in **Figure 4**). Although these TTC tasks differ from the SJ task and were unimodal, both rely on temporal judgments. Therefore, it is plausible that these regions are primarily engaged in temporal processing rather than in multisensory integration per se. It is also possible that they participate in processes common to both tasks, such as attentional and decisional mechanisms mediated by the parieto-frontal [30]. After early sensory processing, auditory and visual cortices interact and converge onto multisensory hubs such as the superior temporal sulcus (STS), which has been consistently implicated as a key site for audiovisual integration and simultaneity perception. From there, frontoparietal regions, including the intraparietal sulcus and frontal areas, likely intervene at a subsequent stage to modulate the sensory signals via top-down mechanisms driven by context, attentional demands, and task goals. These higher-order regions may then feed into decisional and motor circuits, with areas such as the supplementary motor area contributing to the selection and initiation of the behavioral response in the SJ task.

In line with the A Theory of Magnitude (ATOM) framework [53,54], which proposes that time and space may rely on partially shared neural substrates, fMRI work by Skagerlund et al. has identified two regions, in the right intraparietal sulcus and the pre-SMA, (in violet in **Figure 4**) whose activity shows common effects across temporal and spatial magnitude dimensions [55]. This convergence further supports the view that the right parietal and frontal regions highlighted by our DES mapping, together with the right areas reported by Baurès et al. [32,33] constitute a domain-general magnitude network that contributes to temporal and spatiotemporal judgments in both TTC and SJ contexts.

Although DES offers spatial and temporal precision far superior to most other brain mapping techniques, it is not without limitations. First, DES only stimulates cortical surface areas, excluding deeper brain structures. Moreover, operated patients may exhibit tumour-induced cortical reorganization due to neuroplasticity, meaning that regions involved in the task may differ from those typically recruited in a healthy brain. In lesion-based approaches, such reorganization can lead to compensatory activity in regions not originally responsible for the function in question. Given that each patient presents with a unique tumour location and size, group-level statistical analyses are not feasible. Finally, given the constraints of the operating room environment, the number of tasks tested by DES can be rather limited (nevertheless, this is a typical pattern of all brain mapping techniques, invasive or not). As a result, regions considered as ‘task-specific’ may, in fact, be engaged in other functions not assessed intraoperatively. Other brain-mapping techniques, such as transcranial magnetic stimulation (TMS), which are less precise spatially than DES, could serve as a valuable complementary method. TMS can be applied to healthy participants, allowing validation of findings obtained during awake surgery without the same constraints on the number of trials, the stimulated regions, or clinical priorities. For example, TMS was used to investigate temporal-order judgments and demonstrated that modulation of the right posterior parietal cortex (rPPC) widened the TBW [56].

Awake brain surgery, which offers unmatched spatial and temporal resolution, enabled us to causally identify a right parieto-frontal network, encompassing regions near the IPS and SMA, as a key modulator of the audiovisual temporal binding window. Beyond their theoretical contribution, these findings have direct clinical implications by underscoring the need to preserve this network during surgery to avoid postoperative deficits in cognitive functions essential to everyday life.

## Acknowledgments

The authors wish to express their sincere gratitude to the Neurosurgery Department of CHU Purpan for providing access to patients and facilitating the intraoperative recordings. We are also grateful to the surgical teams for their invaluable collaboration.

## Availability of data and materials

Data that supports the findings of this study is available on this link: https://osf.io/ach2n/?view_only=e8673372d24a4e5891c2d5d38d6ee233

## Author contributions

- **Conceptualization:** Solène Leblond, Robin Baurès, Céline Cappe, Franck-Emmanuel Roux
- **Data curation:** Solène Leblond, Robin Baurès, Tutea Atger, Marina Poinsignon
- **Formal analysis:** Solène Leblond, Robin Baurès
- **Investigation:** Solène Leblond, Tutea Atger, Marina Poinsignon, Franck-Emmanuel Roux
- **Methodology:** Solène Leblond, Robin Baurès, Céline Cappe, Franck-Emmanuel Roux
- **Project administration:** Solène Leblond, Robin Baurès, Céline Cappe, Franck-Emmanuel Roux
- **Resources:** Franck-Emmanuel Roux
- **Software:** Solène Leblond (experiment code), Solène Leblond & Robin Baurès (R data analysis scripts)
- **Supervision:** Robin Baurès, Céline Cappe, Franck-Emmanuel Roux
- **Validation:** Solène Leblond, Robin Baurès, Céline Cappe, Franck-Emmanuel Roux
- **Visualization:** Solène Leblond, Robin Baurès, Céline Cappe, Franck-Emmanuel Roux
- **Writing – original draft:** Solène Leblond
- **Writing – review & editing:** Solène Leblond, Robin Baurès, Céline Cappe, Franck-Emmanuel Roux

## Declaration of generative AI and AI-assisted technologies in the writing process

Generative AI and AI-assisted technologies have been used in the writing process to improve the readability and language of the manuscript.

