## Supplementary Material for "Cerebral bases of audiovisual temporal binding window: an awake surgery study"

**<u>Supplementary material 1:</u> Statistical analysis**

We followed the methodology described by Baurès et al. [32, 33]. By contrast with standard experiments, no ANOVA-like analysis can be performed to compare at the group level the influence of the stimulation. Indeed, our patient presented a high degree of heterogeneity in their profile (in particular, tumor location, size, and duration), and consequently, they were not stimulated in the same areas. It was therefore not possible to perform group analysis (e.g., ANOVAs) to determine the influence of the stimulation, as the stimulation differed in localization. We adopted a different strategy, in which each patient served as his own control. We therefore compared, at the patient level, his baseline (pre-surgery) performance with his performance during the surgery condition. For this reason, if age is known to modulate the width of the TBW [29], it cannot explain the variation in a single participant's synchrony perception from the pre-surgery to the per-surgery condition. Note, however, that this method assumes that a participant's pre-surgery performance is stable over time, that is, no practice effects or perceptual learning occur. We specifically controlled this hypothesis by performing test–retest measurements in a control population, as presented in Supplementary Results S1.

To determine whether stimulation of a specific area caused an inability to perform the task correctly, and thus to infer the causal role of the area in the SJ task, we calculated the probability of perceiving the stimuli as synchronous k times out of six repetitions, given that the initial SOA produced 10% or 90% of "synchronous" responses during the pre-surgery condition. This probability was computed using the binomial distribution, described by the following equation:

$$P(X = k) = \binom{n}{k} p^k (1 - p)^{n-k}$$

In which

$$\binom{n}{k} = \frac{n!}{k!\,(n-k)!}$$

For example, for a 10% probability of "synchronous" responses ($SOA_{10}$), the probability p to perceive the stimuli as synchronous k = 0 times over the 6 repetitions is *p* = .531, p = .354 when k = 1, *p* = .098 when k = 2, p = .015 when k = 3, p = .001 when k = 4, p = 5.4e−05 when k = 5, and p = 1e−06 when k = 6.

Since $SOA_{10}$ corresponds to a large delay, perceiving auditory and visual stimuli as asynchronous should be the norm, making a high number of "synchronous" responses unexpected. Accordingly, we considered it unlikely that perceiving the $SOA_{10}$ as synchronous three or more times (cumulative probability of 3 or more is p = .016) would indicate involvement of the deactivated area in the SJ task. Symmetrically, for $SOA_{90}$, the threshold was set at perceiving the stimuli as synchronous three times or fewer to conclude the area's involvement in the perception.

**<u>Supplementary material 2:</u> Table of stimulation coordinates (MNI coordinates) and left brain region localization of patients tested in awake surgery**

| Patient number | X (MNI) | Y (MNI) | Z (MNI) | Effect of stimulation | Localization |
|---|---|---|---|---|---|
| 1 | -62,4 | -16,8 | -10,9 | No effect | Middle temporal gyrus |
| 2 | -37,7 | 27,9 | 44,2 | No effect | Middle frontal gyrus |
| | -31,1 | 33,1 | 45,1 | No effect | Superior frontal sulcus |
| 3 | -32,3 | 15,3 | 55,7 | No effect | Superior frontal sulcus |
| 4 | -52,1 | 0,5 | -24,7 | No effect | Middle temporal pole gyrus |
| | -57,7 | -2,8 | -18,2 | No effect | Middle temporal gyrus |
| | -55,1 | 18,2 | 7,6 | No effect | Inferior frontal gyrus |
| 5 | -61,8 | -28,5 | 10,6 | No effect | Superior temporal gyrus |
| | -61,1 | -25,6 | 19,8 | No effect | Supramarginal gyrus |
| 6 | -60,5 | -35,2 | 13,2 | No effect | Superior temporal gyrus |
| | -60,2 | -24,4 | 29,1 | No effect | Supramarginal gyrus |
| 7 | -29,3 | 28,7 | 49,7 | No effect | Superior frontal sulcus |
| 9 | -61,5 | -43,4 | 11,1 | No effect | Superior temporal gyrus |
| | -59,8 | -47 | 17,5 | No effect | Superior temporal sulcus |
| 10 | -50,3 | 3,9 | -23,4 | No effect | Middle temporal pole gyrus |
| | -52,4 | 8,6 | -9,3 | No effect | Superior temporal gyrus |
| | -62,5 | -21,7 | 6,7 | No effect | Superior temporal gyrus |
| 11 | -30,8 | 49,7 | 31,5 | No effect | Middle frontal gyrus |
| | -32,8 | 45,6 | 33,5 | No effect | Middle frontal gyrus |
| | -36,4 | 34,6 | 39 | No effect | Middle frontal gyrus |
| 12 | -52,3 | 35,4 | 4,2 | Language + SJ | Inferior frontal gyrus |
| | -49,8 | 40,1 | 9,6 | No effect | Inferior frontal sulcus |
| 13 | -49,6 | 12,7 | -15,8 | No effect | Superior temporal gyrus |
| 14 | -39,5 | -79,6 | -0,8 | No effect | Lateral occipital gyrus |
| 15 | -37,5 | 12,4 | 50,5 | No effect | Middle frontal gyrus |
| | -32,6 | 7,8 | 56 | No effect | Superior frontal sulcus |
| 18 | -56,8 | -1,9 | -1,8 | No effect | Superior temporal gyrus |
| 19 | -41 | -64,5 | 39,3 | No effect | Angular gyrus |
| 20 | -52,7 | 26,6 | 20,6 | No effect | Inferior frontal gyrus |
| | -62,6 | -14,6 | -9 | No effect | Middle temporal gyrus |
| 21 | -31 | -55,4 | 54,2 | No effect | Intraparietal sulcus |
| | -40,6 | -51,1 | 48,2 | No effect | Intraparietal sulcus |
| 22 | -46,2 | -53,7 | 44 | No effect | Angular gyrus |
| | -31,3 | -51,8 | 56,5 | No effect | Intraparietal sulcus |
| 29 | -25 | -0,4 | 63,1 | Motor + SJ | Superior frontal sulcus |
| | -9,2 | 0,4 | 67,6 | Language + SJ | Supplementary motor area |
| 30 | -7,5 | 10 | 65,1 | No effect | Supplementary motor area |
| 31 | -58,4 | 10,3 | 21,5 | No effect | Precentral sulcus |
| 34 | -59,2 | -6,8 | -2 | No effect | Superior temporal gyrus |
| 35 | -62,3 | -27,7 | 10,1 | Language + SJ | Superior temporal gyrus |

| | | | | | |
|---|---|---|---|---|---|
| | -66,5 | -29 | 2,5 | Language + SJ | Middle temporal gyrus |
| 36 | -41 | -48,6 | 46,8 | No effect | Intraparietal sulcus |
| | -32,7 | -55,1 | 52,4 | No effect | Intraparietal sulcus |
| 37 | -50,3 | 15,8 | 38 | No effect | Inferior frontal sulcus |

**Supplementary material 3: Table of stimulation coordinates (MNI coordinates) and right brain region localization of patients tested in awake surgery**

| Patient number | X (MNI) | Y (MNI) | Z (MNI) | Stimulation effect | Localization |
|---|---|---|---|---|---|
| 8 | 62,8 | -39,8 | 12,8 | No effect | Middle temporal gyrus |
| | 61,5 | -48,1 | 8,1 | No effect | Middle temporal gyrus |
| 16 | 32,7 | 4,8 | 57,3 | No effect | Middle frontal gyrus |
| 17 | 58 | -37,3 | 36,5 | **SJ-specific** | Supramarginal gyrus |
| 23 | 23,4 | 7,4 | 63,4 | No effect | Superior frontal sulcus |
| | 33 | 8,2 | 56,5 | No effect | Middle frontal gyrus |
| 24 | 10,8 | 7,3 | 66,6 | No effect | Middle frontal gyrus |
| | 15,6 | -0,5 | 68,4 | **SJ-specific** | Supplementary motor area |
| 25 | 38,1 | -50,5 | 55,9 | **SJ-specific** | Intraparietal sulcus |
| 26 | 37,8 | 19,8 | 48,6 | **SJ-specific** | Middle frontal gyrus |
| 27 | 49,9 | -50,4 | 44 | **SJ-specific** | Intraparietal sulcus |
| 28 | 27,3 | -4,8 | 64,4 | No effect | Superior frontal gyrus |
| 32 | 46,2 | 2,1 | 48,9 | No effect | Precentral sulcus |
| 33 | 11,2 | 5,2 | 68 | **SJ-specific** | Supplementary motor area |
| | 10,1 | 13,2 | 64,5 | **SJ-specific** | Supplementary motor area |
| 38 | 35,3 | 2,2 | 58 | No effect | Middle frontal gyrus |
| | 25,8 | 1,3 | 63,8 | No effect | Middle frontal gyrus |
| 39 | 39,1 | -49,3 | 56,2 | No effect | Superior parietal gyrus |
